# Automated Neuron Tracing with Imitation Learning

**DOI:** 10.64898/2026.07.29.741595

**Authors:** Bryson Gray, Daniel Tward

## Abstract

Reconstructing neuronal morphology from large 3D microscopy volumes is essential for quantitative neuroscience but remains challenging due to noise, low contrast, and complex branching geometry. We present an automated neuron tracing method that formulates reconstruction as a sequential decision-making problem and trains a 3D convolutional policy via imitation learning. The key technical contribution is an online expert-action retrieval scheme that derives locally valid continuation directions directly from a gold-standard SWC tree, including at bifurcations. Combined with DAgger-style dataset aggregation, the policy is trained under its own state distribution without requiring handcrafted reward functions or a separate segmentation-to-skeleton postprocessing stage. We package the approach into an end-to-end tracing pipeline with optional path refinement operations and tools for quantitative evaluation and interactive editing. On the BigNeuron Gold166 benchmark, our method achieves state-of-the-art reconstruction accuracy, outperforming all competing challenge methods under both morphology-based aggregate metrics and geometric distance comparisons.

## 1. Introduction

Reconstructing neuronal morphology from 3D microscopy volumes is a vital step in computational neuroscience, enabling quantitative analysis of neural circuits, connectivity, and cell-type classification. Advances in imaging modalities such as confocal microscopy, two-photon microscopy, and serial electron microscopy have made it possible to acquire large-scale volumetric datasets at submicron resolution. However, extracting accurate neuronal reconstructions from these datasets remains a major computational challenge due to imaging noise, low contrast, complex arborization patterns, and the presence of dense, overlapping neurites Parekh and Ascoli (2015); Ascoli et al. (2007).

Classical approaches to neuron tracing typically formulate reconstruction as a global optimization or graph extraction problem. Early methods rely on image filtering and shortest-path algorithms, where tubularity-enhanced images are used to construct graphs and paths are recovered via minimal-cost routing Cai et al. (2006); Peng (2011). Probabilistic formulations, including particle filtering and Bayesian tracing, extend this paradigm by incorporating uncertainty during sequential tracing Zhang (2010). While these approaches are effective in relatively clean datasets, they often struggle with discontinuities, ambiguous crossings, and large-scale branching structures.

More recently, deep learning has significantly advanced the field by enabling data-driven feature extraction. Convolutional neural networks (CNNs) and 3D U-Net architectures have been widely adopted for voxel-wise segmentation of neurites, followed by skeletonization or graph reconstruction Ronneberger et al. (2015); Li (2017); Banerjee et al. (2025). These methods achieve strong performance, particularly when large annotated datasets are available. However, segmentation-based pipelines typically decouple local appearance modeling from global topology reconstruction, which can lead to topological errors such as broken branches, false merges, or spurious loops. Moreover, converting dense segmentations into accurate tree structures remains a non-trivial postprocessing step. An alternative perspective is to formulate neuron tracing as a sequential decision-making problem, where a tracer iteratively extends a neurite by selecting its next direction conditioned on local image context. This view aligns naturally with reinforcement learning (RL), and previous works have explored learning navigation policies for neurite tracing in 2D microscopy images Dai (2019); Balaram et al. (2019). Although RL-based approaches are promising, designing a reward function that can produce the desired tracing behavior in three-dimensional image volumes, including correct branching and termination of paths is a challenge. As a result, their adoption has remained limited compared to supervised learning approaches. Furthermore, RL was designed for problems where the inner workings of an environment can only be examined by taking actions, and where optimal actions are unknown. By design, they do not make use of all data available for neuron tracing tasks.

In contrast, imitation learning (IL) provides a compelling framework for leveraging expert knowledge in sequential decision problems. Instead of relying on manually designed reward functions, IL enables a model to directly learn tracing policies from expert demonstrations, such as manually curated neuronal reconstructions. In particular, dataset aggregation (DAgger) offers a principled approach to mitigate compounding errors by iteratively collecting training data under the learned policy while querying an expert for corrective actions Ross et al. (2011). Despite its success in robotics and autonomous navigation, imitation learning has been relatively underexplored in the context of neuron reconstruction. A key challenge in applying imitation learning to neuron tracing lies in defining expert actions in a continuous, branching 3D environment. Unlike standard navigation tasks, neuronal structures exhibit complex geometry, including tortuous paths, varying radii, and frequent bifurcations. Existing approaches often rely on local heuristics or discretized direction sets, which may not capture the underlying morphology. This motivates the need for principled methods that derive supervision signals directly from expert reconstructions while respecting the geometric and topological properties of neuronal trees. In this work, we present NeuroTrack, a novel imitation learning framework for 3D neuron tracing that leverages expert morphological reconstructions to define target tracing steps. Our method introduces a new target step estimation strategy that extracts locally optimal continuation directions from ground-truth neuron skeletons, enabling accurate supervision of the tracing policy even in the presence of branching. Combined with a DAgger-based training procedure, our approach iteratively refines the policy to remain robust under its own state distribution, reducing error accumulation during long-range tracing.

We develop and package our tracing method along with postprocessing and evaluation tools to provide a full pipeline for tracing, refinement, and evaluation and comparison with existing methods. We also provide a graphical user interface along with our command-line tool to enable easy selection of seed points, visualization of traces, and manual editing of automated traces. Finally, we apply our method to the BigNeuron Gold166 challenge dataset to directly compare our method to 34 other state-of-the-art neuron tracing algorithms. Our tracing, processing, and evaluation pipeline is illustrated in Figure 1.

**Figure 1:**
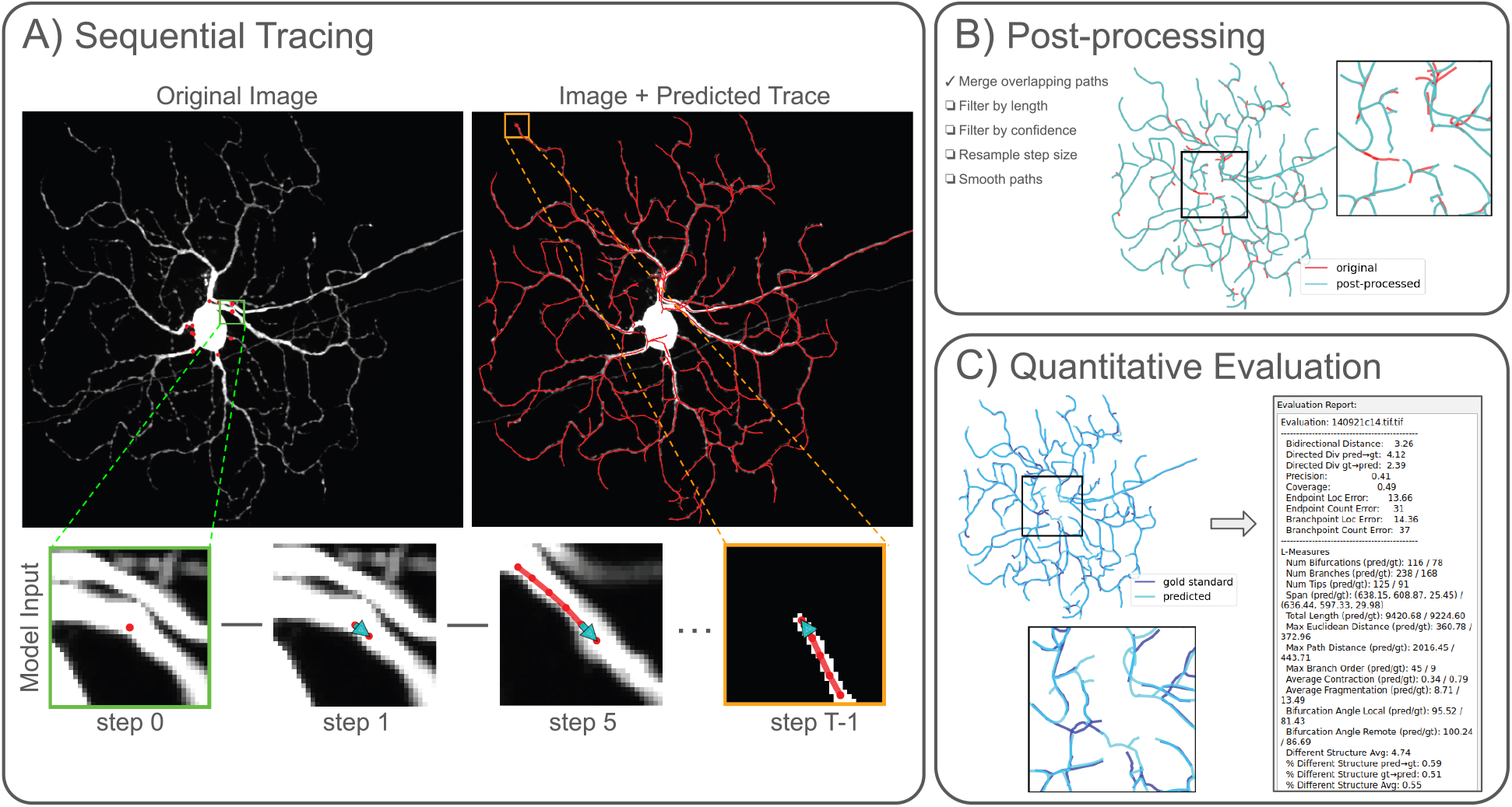
A) Automated sequential tracing procedure. Each trace begins at a provided seed point and proceeds stepwise until all paths have terminated. The agent (a 3D CNN) takes local image patches as input and outputs step directions. B) Paths may be refined by several optional postprocessing operations: Merge overlapping paths, filter by length, filter by confidence, resample step size, and smooth paths. C) Quantitative reconstruction evaluation was performed by comparing the reconstruction with gold standard annotations. Evaluation metrics are described in section 2.7.

We hypothesize that framing neuron reconstruction as an imitation learning problem offers several advantages: (1) it naturally incorporates expert knowledge without requiring handcrafted reward functions, (2) it enables stable training compared to RL-based methods, and (3) it optimizes sequential tracing performance directly rather than intermediate segmentation quality. Our empirical results show that the proposed method outperforms all other methods included in the Gold166 challenge when comparing both an aggregate metric of morphological measures and geometric distance alone in direct match-ups, lending support to these hypotheses and highlighting imitation learning as a promising direction for scalable, high-fidelity neuron reconstruction. To our knowledge, this is the first work to apply DAgger-style imitation learning to neuron tracing.

## 2. Methods

### 2.1. Data

Data for training and evaluation were obtained from the BigNeuron Gold166 challenge dataset Manubens-Gil et al. (2023), which comprises 166 expert-annotated neuronal reconstructions in SWC format paired with 3D microscopy image volumes. Several volumes were excluded because their corresponding gold-standard reconstructions did not reliably follow the neurite centerlines evident in the image data and could not be corrected by applying a uniform spatial offset. Additional volumes were excluded because they contained binary intensity values rather than raw image intensities, resulting in insufficient contrast in densely arborized neurite regions. After this filtering step, 134 image volumes remained. We refer to this dataset as a “gold-standard” throughout. Its original authors have acknowledged potential limitations (some reconstructions can be ambiguous in cases of limited image quality) but described this dataset as “the best possible reconstruction generated by human experts from the respective image given the limited image quality, time and resources”.

Our method for extracting optimal supervision targets requires that the reference neuron reconstruction is an acyclic, connected graph (i.e., a tree). However, many of the gold-standard reconstructions included disconnected components, duplicate node identifiers, and nodes occupying identical world coordinates. As a result, these reconstructions required additional preprocessing before they could be used as training targets. Our training algorithm relies on the gold standard annotations to compute expert step directions, but many neurons have large somas inside which there is no clear “correct” step direction. Therefore, to prevent the annotated regions inside the somas from adding noise to the ground truth labels, we use the gold standard annotations with the soma removed for both training and testing annotations. Somas were removed using our graphical interface by drawing a bounding box around the soma in the neuron image and removing the gold standard annotation nodes that fall inside the bounding box. The preserved nodes nearest to the bounding box boundary were stored as seed positions for path initialization. These were used for both training and testing to help reduce bias in seed point selection. Finally, all retained images were cropped to the minimum and maximum neuron coordinates, and the coordinates were translated accordingly, in order to decrease training time by reducing the cost of loading data into memory.

Supplementary Table 3 lists the neurons and whether each was included or excluded from our dataset. More information about our inclusion/exclusion criteria and preprocessing steps are detailed in the dataset filtering and preprocessing subsection in Supplementary Information.

### 2.2. Environment

The neuron tracing environment is what the agent (a 3D CNN that learns a policy) interacts with during training. It records the paths traced by the agent, increments the current position, returns the next state observation given the agent’s action, and computes the expert target steps from the expert-annotated neuron tree during training. The environment also resets when the trace is complete, ending the episode and loading the next image to be traced. Each training episode includes a neuron image volume together with its corresponding gold standard neuron tree, a binary neuron mask (obtained by binarizing voxels based on their distance to the neuron tree with a cutoff of 17 voxels) used to govern path termination, and a set of seed points for path initialization. During inference, only the image volume and seed points are given. The agent takes a local observation around the current path head as input. The observation is a two-channel sub-volume with shape 2 x 35 x 35 x 35 where the channels are (1) the image and (2) a path-history channel showing the path history as a binary mask with ones along the predicted path. The path-history channel is necessary to remove the ambiguity as to which direction along the neuron to proceed (i.e. the agent should proceed in the direction that has not yet been traced). Given a local observation, the agent outputs an action as a 3D vector which is added to the previous path position to advance the path. At each step, the environment categorizes the action as stop or continue depending on the length of the vector (i.e. step sizes smaller than one voxel are treated as a stop signal). The environment maintains a list of positions at each step: a neuron tree which will ultimately be converted to a SWC file. And it produces a set of candidate branch roots (Figure 2 green circle), which are added to a first-in-first-out queue as the agent progresses. The agent will return to each candidate branch root, which may be extended into a longer path if appropriate.

**Figure 2:**
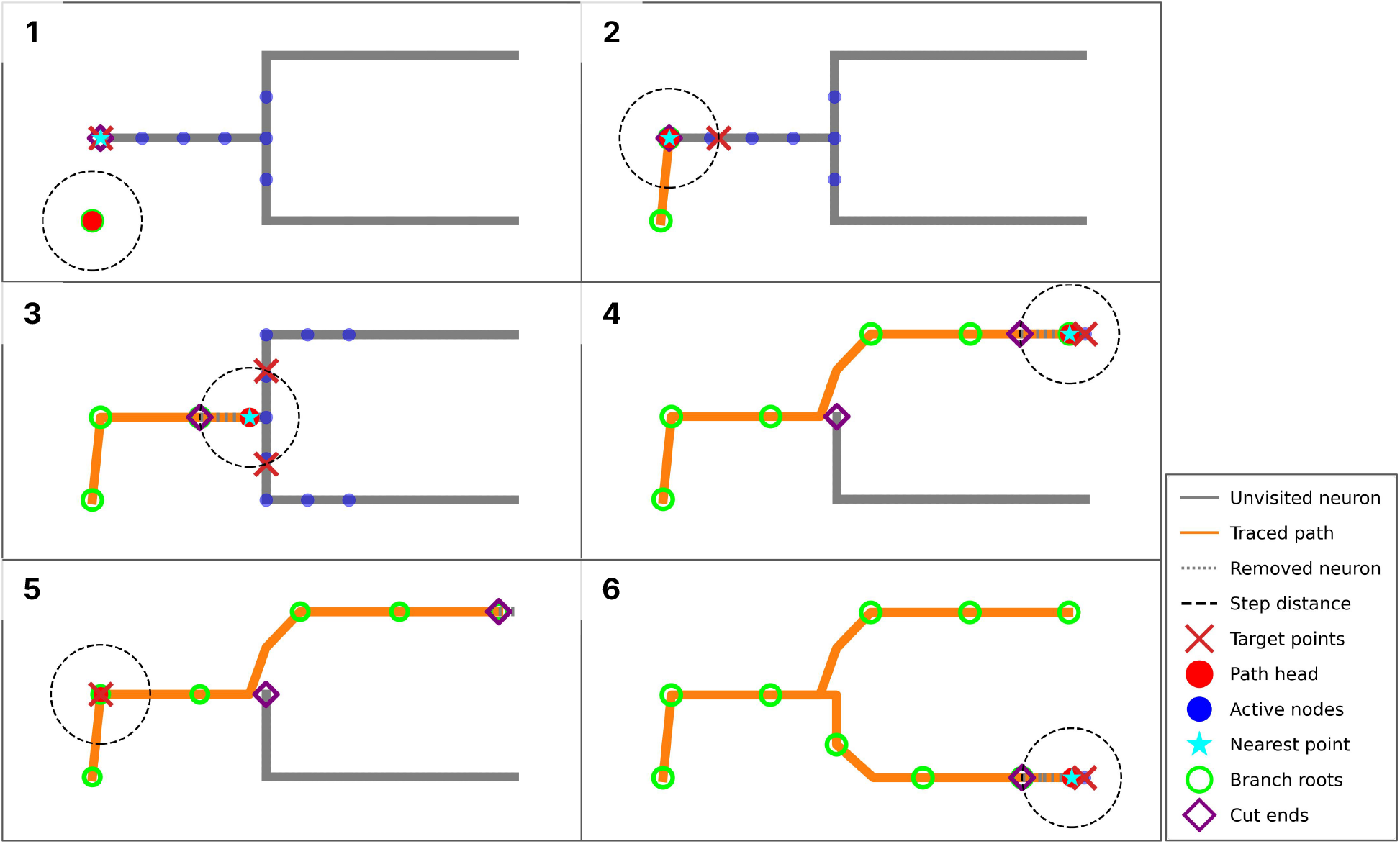
Expert action retrieval from reference SWC and traversed edge bookkeeping. The neuron tree is removed as the path (orange line) progresses along the neuron (gray line), always removing sections *from* a cut end (purple diamond) *to* the nearest point to the new position (cyan star). The section of neuron tree removed in the last step is shown as a dashed gray line. New branch roots (green circle) are automatically placed at regular intervals and stored in a first-in-first-out queue. These branch roots will be revisited, and may grow into branches. (1) The path begins at the first branch root. No neuron point lies at the step distance (broken black line) so the target point (red ‘X’) is placed at the nearest neuron point. (2) The agent takes the first target step. (3) Near branch points multiple neuron points lie a step distance from the path head. The nearest candidate to the step taken is used as the training target. (4) When a terminal is less than the step distance away, it becomes the target. The path stops near the terminal as a consequence of taking a step shorter than the stop threshold distance. (5) When the first branch terminates, the current path moves to the next branch root in the queue. The new branch shown is farther than a threshold distance from any valid nodes so the target step is a zero vector which tells the environment to terminate the path. (6) Tracing continues from the next queued branch until all paths have terminated.

**Figure 3:**
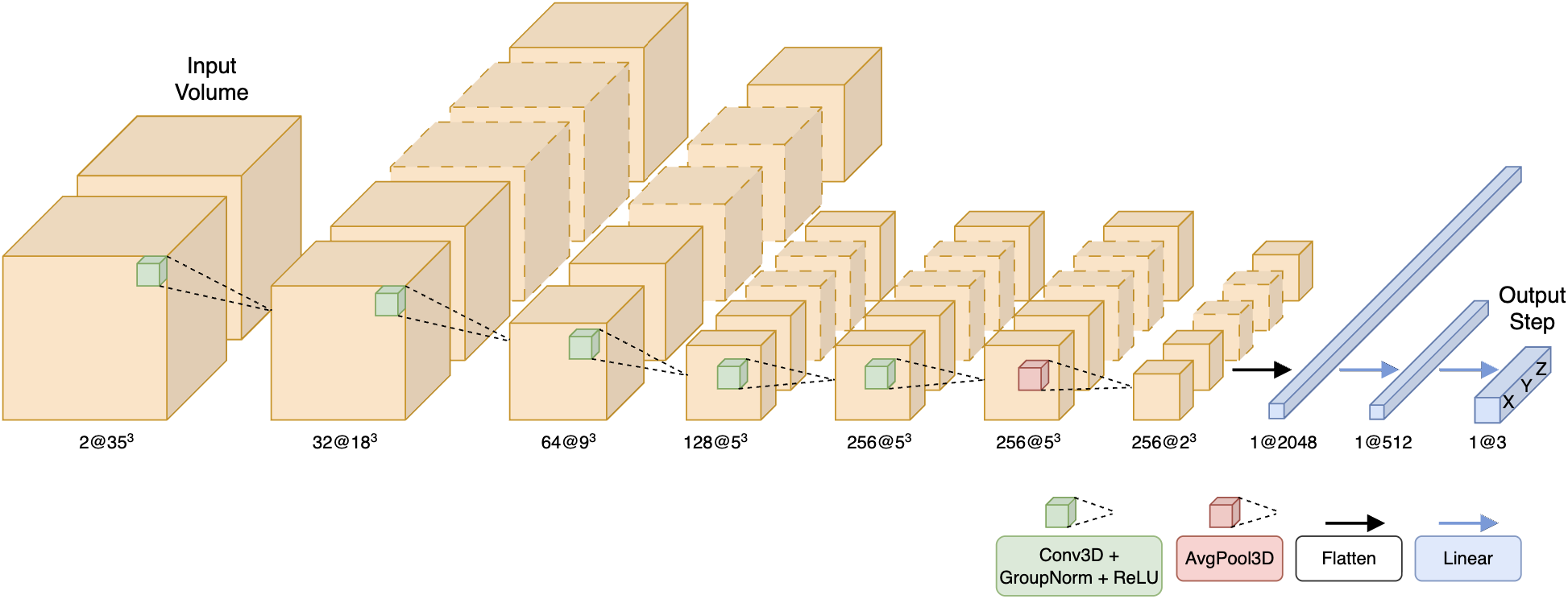
3D Convolutional Neural Network Architecture: Five convolutional layers, each followed by a GroupNorm and ReLU operation. The first three convolutional layers use a stride of two for downsampling. All convolutions have a kernel size of three and padding set to one. The convolution layers are followed by average pooling, flattening to one dimension, and two linear layers resulting in a 1×3 output step vector.

Our environment uses an online supervision strategy that algorithmically derives expert actions directly from the evolving neuron tree during tracing. As a result, no matter where the learning agent moves within the image space, the true next action is computed on the fly and supplied as a training label, eliminating the need for human intervention (in contrast to certain DAgger applications that require human annotators Kelly et al. (2019); Han et al. (2026); Xu et al. (2025)). Figure 2 provides an idealized illustration of the interaction with the environment. Given the agent’s current position (Figure 2 red dot), one or more target points are computed by finding all points on the neuron that are a specified step distance away (5.0 voxels in our configuration) from the current path position, noting that such points are generally on line segments between SWC nodes rather than the nodes themselves (Figure 2 red ‘X’). The target points are then converted to relative direction vectors. Near branch points, multiple valid candidates are preserved (Figure 2 panel 3). The training target is the candidate direction closest to the predicted direction, which yields supervision targets that flexibly handle the topology of the branching neurons.

It may be that no untraced neuron points exist at exactly the step distance. This will occur either when the agent approaches a terminal point (Figure 2 panels 4 and 6), when it is located at a candidate branch root that does not correspond to a real bifurcation (Figure 2 panel 5), or when the agent is far from the neuron, perhaps due to a poor seed point or erroneous step (Figure 2 panel 1). In these cases, the expert action steps to the neuron point nearest to the step distance on the connected component nearest to the current position (Figure 2 panel 2), provided it is nearer than a predefined cutoff distance. Following this logic, when the path approaches a true termination point to less than the step distance, the expert action points to that terminal, and once the distance drops below the stop threshold of one voxel, triggers the path to terminate (Figure 2 panels 4 and 6). If it is larger than the cutoff distance, the expert action is assigned zero length, which is interpreted by the environment as a signal to terminate the path (Figure 2 panel 5).

To prevent failure modes common in tracing, we explicitly remove the already traversed portions of the gold-standard tree as the agent moves (Figure 2 Note that gray is replaced by orange). The fraction of each edge that has been traversed is recorded. Fully traversed edges are deleted from the working tree, and partially traversed edges are split at “cut ends”. At each step, we mark as “traversed” and remove from the working tree all edges along the neuron between the cut end closest to the start position and the active neuron point (a point in the “active region”, defined below) nearest to the new position. This ensures that subsequent expert targets are sampled only from the remaining unvisited parts of the tree, enforcing forward progress.

While taking a step, both the target points and the start/end points for edge removal are restricted to a local region of the working tree, the active region, indicated in Figure 2 by blue dots, defined as the set of nodes within a bounded geodesic distance (i.e. distance measured by walking along a tree, rather than straight through the background space) of any nearby cut end created by previous edge removals. This restriction prevents expert targets from crossing between neighboring branches and taking shortcuts across sharp turns.

Figure 2 panels 1 to 6 describe a sequence of steps taken during a typical episode. The details, which refer to the components discussed in this section above, are enumerated in the figure caption.

### 2.3. Dataset Aggregation Training Algorithm

Behavior cloning, considered the most basic formulation of imitation learning, trains a policy using a fixed training set consisting of states paired with expert actions. This often fails in practice because in sequential prediction problems, each state depends on each previous state, violating the i.i.d condition often assumed in statistical learning Ross et al. (2011). Consequently, the distribution of states induced by the trained policy at deployment time may differ substantially from the states represented in the training data, leading to cascading deviations from expert behavior. The Dataset Aggregation (DAgger) algorithm mitigates this issue by iteratively collecting data from the learner’s own state distribution while querying an expert for corrective labels. Concretely, DAgger alternates between executing the current policy to induce a trajectory distribution and aggregating expert-labeled state–action pairs into a growing dataset, on which the policy is repeatedly retrained. This process improves robustness by ensuring coverage of off-policy states encountered during learning.

Our formulation of the DAgger algorithm begins with a warm-start phase in which the model learns exclusively on the expert trajectories. During this phase, each environment step stores the observation and the full set of candidate target vectors, and a deterministic expert action is selected from the candidates to advance to the next state (favoring alignment with the previous action when multiple targets exist). Data collection continues until the number of steps taken meets the designated steps per update, *T*_update_, optimization is performed, and then data collection resumed from where it paused. The warm-start phase ends when the total steps taken meets the warm-start length, *T*_ws_.

After the warm-start phase, a new data collection round begins (a “roll-in round”). This time we begin to roll in actions from the learned policy at random with probability *β*. Given the policy learned by our agent 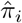 and expert policy *π*^*∗*^, the policy at round *k* is 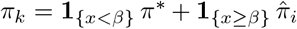 where *x ~ U* (0, 1). Regardless of which policy the action is sampled from to advance to the next state, the expert is always queried for the target vectors to add to memory. Every *T*_*update*_ steps, the collected state-action pairs are added to the training dataset and the policy is updated by optimizing over the aggregated dataset. Collection resumes after optimization and runs until the total steps collected in the round reaches the specified steps per DAgger round, *T*_DAgg_. For the next collection round *β* may be reduced, increasing the frequency of on-policy actions and potentially out-of-distribution states. After each optimization, we evaluate the learned policy by using it to trace all the images in the training image set and computing the average symmetric distance between the predicted and gold standard reconstructions. If the new average distance is smaller than the smallest achieved so far, then we reduce beta by a fixed increment, *β*_step_. The gradual reduction of *β* during the course of training prevents the policy from trying to learn behavior for states that are not likely to be relevant later. The algorithm is detailed in Algorithm 1.

#### Algorithm 1

DAgger training procedure with an initial warm-start phase using the expert policy before gradually mixing in the learned policy and adapting the mixing coefficient based on evaluation performance.

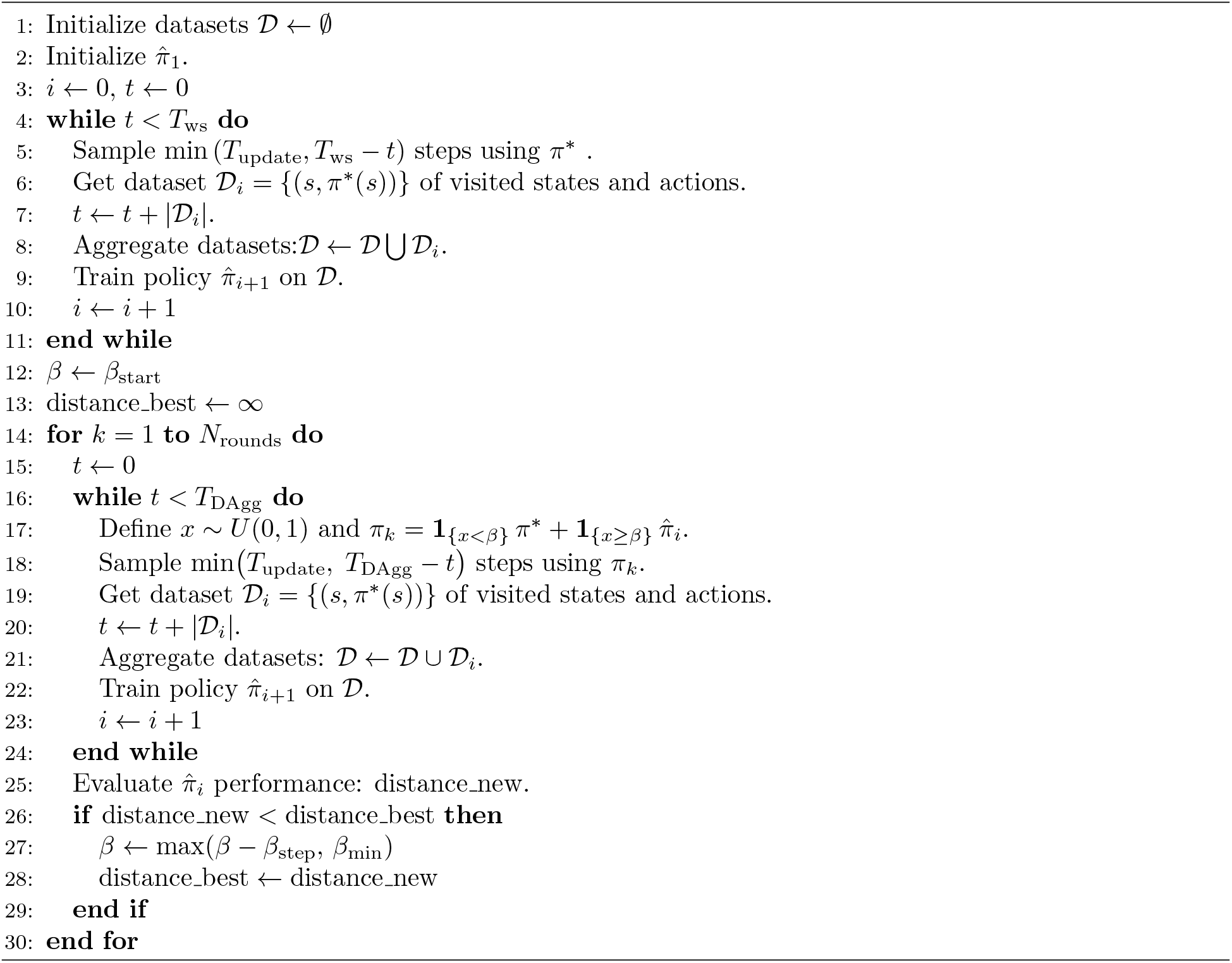

### 2.4. Policy Optimization and Memory Buffer

Policy optimization is done by sampling batches from the aggregate memory buffer and minimizing a multi-target direction loss: for each batch, the loss is the average squared distance to the nearest valid target vector (see Fig. 2c) plus binary cross-entropy (BCE) stop/continue classification.

Let 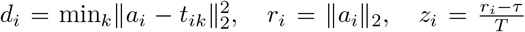, *C* = *{i* : *r*_*i*_ *> τ}, S* = *{i* : *r*_*i*_ *≤ τ}*, where *a*_*i*_ is the predicted action at step *i, t*_*ik*_ are target candidates, *τ* is the continue-norm threshold, and *T* is the stop-classification temperature.

#### Direction loss

For points in the continue set *C*, the direction loss is

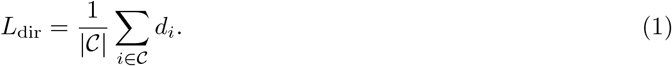

#### Classification loss

We define positive (continue) and negative (stop) classification losses as

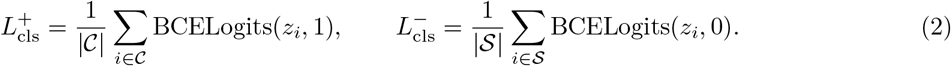

The combined classification loss is then

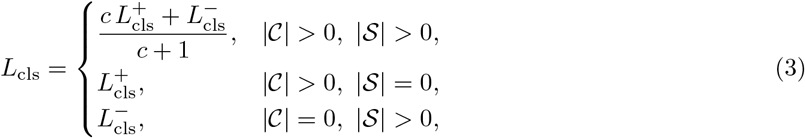

where *c* is the continue weight.

#### Final loss

The overall training objective is

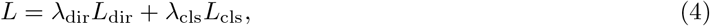

where *λ*_dir_ is the continue-direction weight and, *λ*_cls_ is the norm-classification weight.

During each policy update step, one training epoch is run over the aggregated memory buffer. Data augmentation is applied at sampling time by randomly permuting spatial axes (swapping the two in-plane axes) and randomly flipping each axis. This augmentation is applied consistently to the observation and the target vectors. Optimization is performed using the AdamW algorithm.

The aggregate memory buffer has a fixed capacity. Once it reaches capacity, new samples randomly overwrite old samples. This random replacement of samples was chosen over a straight-forward first-in-first-out rule in order to prevent the oldest samples from being replaced too quickly.

### 2.5. Neural Network Architecture

The policy is a 3D convolutional network that maps a local image patch to a 3D action vector. It consists of five Conv3D blocks, each followed by GroupNorm normalization and a ReLU layer. A global AdaptiveAvgPool3d reduces the feature map dimensions, which is flattened and passed through two fully connected layers with LayerNorm and ReLU on the hidden layer. We use Kaiming (He) initialization for convolutional and linear weights He et al. (2015). We instantiate the model with two input channels (image plus path-history channel) and three output channels to predict a 3D direction vector, with observations normalized from uint8 to [0, 1] before inference.

### 2.6. Trace initialization

Tracing must start from one or more designated seed locations. For model training, validation, and testing, we rely on near-soma seeds obtained from the gold standard annotations, as detailed in section 2.1. During model validation, we found that small perturbations in seed locations could significantly alter the final trace and the resulting evaluation metrics, primarily due to occasional stop-versus-continue misclassifications. To improve robustness against this type of error, we introduced an optional random seed jittering procedure, in which each provided seed point generates *N*_jitter_ additional seeds by randomly translating its position inside a specified radius. This strategy is similar to tractography approaches used in analysis of diffusion MRI (Behrens et al. (2007); Tournier et al. (2012); Correia et al. (2008)). Instead of uniformly jittering the seeds, we use boundary-weighted random sampling, where a Sobel filter is applied to the seed region to generate an image gradient map that then serves as the sampling weights. When multiple seed points are provided, the path-history channel is cleared after completing the trace from one seed and before beginning from the next seed. This prevents traces rooted at nearby seeds from affecting each other. However, this strategy produces traces that are mostly redundant: multiple paths trace the same neurite, raising the need for a postprocessing step to merge redundant paths. It also creates an opportunity: if we consider the number of paths within a vicinity to approximate the confidence that they trace a true neurite, we can use this number to filter out low-confidence traces, which are more likely to be false positives. The postprocessing steps are detailed in the next section.

### 2.7. Postprocessing

Our automated tracing algorithm is biased toward over-tracing for two reasons: (1) Because the agent revisits the length of each trace in discrete intervals to repeatedly evaluate whether a genuine branch is present, it can generate short false-positive branches when it takes one or two small steps before stopping.

(2) When random seed jittering is turned on, many paths redundantly trace the same neurites. Even with seed jittering turned off, we occasionally observed separate trees originating from nearby seed locations such that one seed captured both the proximal dendrite and the dendrite nearer to the neighboring seed. To mitigate these problems, we incorporated several postprocessing steps into our neuron tracing pipeline. Each step is optional and may be included or omitted depending on the specific use case.

The postprocessing operations are as follows: (1) Confidence clipping: When seed jittering is enabled, the density of traced paths may be used to estimate the confidence that a true neurite is present. Confidence clipping first maps the number of nearby paths for each node in the full reconstruction, then finds long runs of consecutive low-confidence nodes, defined by a threshold number of nearby paths (confidence threshold), and clips the tree at the beginning of every low-confidence run, removing the clipped node and all of its descendants. (2) Merging: First, an overlap mask is created, identifying all node indices which are within a threshold distance from a longer path (merge threshold). Where a path overlaps with a longer path, the nodes are removed. If the overlap region is followed by a non-overlapping region, the start of the non-overlapping run is re-anchored to the nearest node on the path that it overlapped. (3) Length filtering: paths are filtered by length. Only paths whose lengths are within a user-defined length range are retained. When a path is removed, all its descendants are removed to avoid creating disconnected sections. (4) Resampling: every segment of the tree, defined as a connected sequence of nodes bounded by either end points or branch points, is resampled to a regular step interval. The boundary points are preserved to avoid changes to the topology. (5) Smoothing: The traced paths are smoothed using a moving average of the path coordinates.

### 2.8. Evaluation

Reconstruction quality was assessed with a suite of metrics combining geometric agreement and morphology statistics. First, we measure the average nearest-neighbor distance in both directions, from the reconstructed tree to the reference tree and from the reference tree to the reconstruction, and use their mean as an overall symmetric distance error. Using a fixed spatial tolerance (2 voxels), we then quantify how many points in each direction are not sufficiently matched; from these mismatch fractions, we report one score reflecting how much of the reconstruction is supported by the reference (precision) and one score reflecting how much of the reference is recovered by the reconstruction (coverage). To evaluate topological landmarks, we identify terminal nodes and branching nodes in each tree and compute the symmetric localization error for each landmark type, along with absolute differences in their counts. We also report L-measures Scorcioni et al. (2008) for both reconstruction and reference: number of bifurcations, number of branches, number of terminal tips, spatial extent along each axis, total cable length, maximum straight-line distance from the root, maximum path length from the root, highest branch order, mean contraction (straight-line-to-path ratio), mean branch fragmentation, and local and remote bifurcation angles. Finally, we summarize structural disagreement beyond the distance tolerance by reporting the mean distance among only mismatched points (different structure average) and the directional and averaged proportions of mismatched points (different structure fraction). The metrics are summarized as follows:

We first define *D*_*G*_(*x*) as the distance of a point *x* to the nearest point on neuron tree *G*:

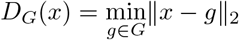

- **Directed geometric distance**. The mean nearest-neighbor distance from each point in tree *A* to tree *B* is

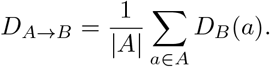 Lower values indicate better geometric agreement.
- **Symmetric geometric distance**. We report the symmetric average of the two directed distances:

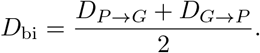 Here *P* is the predicted reconstruction and *G* is the ground truth neuron tree.
- **Different structure average**. Using a tolerance *τ*, we define

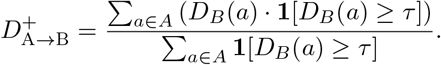

i.e. the average distance of points in *A* to *B* for those that are farther than *τ*. We report the symmetric different structure average:

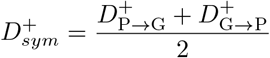
- **Different structure fraction**. We report the fraction of points exceeding *τ* in each direction and their average:

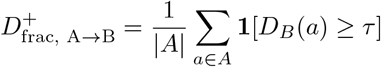

and the symmetric different structure fraction:

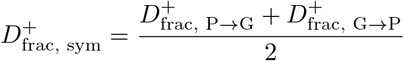
- **Precision**. Using a tolerance *τ*, we define

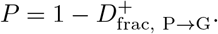 Higher values indicate better support of the reconstruction points by the reference neuron.
- **Coverage**. Using the same tolerance *τ*, we define

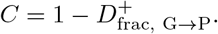 Higher values indicate better recovery of the reference.
- **Endpoint localization error**. We detect terminal nodes in each tree and report symmetric nearest-neighbor localization error:

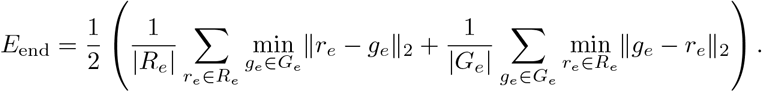 Lower values are better.
- **Branchpoint localization error**. We compute the same symmetric nearest-neighbor error for branchpoints:

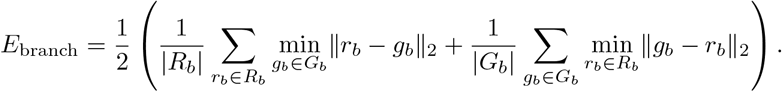
- **Endpoint count error**. We report the absolute difference in the number of terminal nodes:

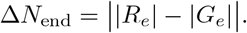
- **Branchpoint count error**. We report the absolute difference in the number of branch nodes:

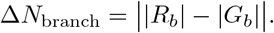
- **Other morphology measures**. We also report reconstruction and reference morphology separately using: number of bifurcations, number of branch segments, number of terminal tips, spatial span along each axis, total path length, maximum Euclidean distance to the root, maximum path distance to the root, maximum branch order, average contraction, average fragmentation, and local and remote bifurcation angles.

### 2.9. Experiments

Our model was trained using a 5-fold cross-validation scheme. Because the full dataset is composed of thirteen subsets, each originating from a different laboratory and/or subject species, we balanced the folds by ensuring that every split contained approximately equal proportions of images from each subset (See Supplementary Table 4 for fold information). The model was trained for 100 roll-in rounds while saving model parameter checkpoints after each round. The checkpoint that scored the best on in-training evaluation (see section 2.3) was selected for the final model.

The trained model reconstructions were produced for the held-out test fold under two conditions: (i) using only seed points derived from the gold-standard annotations, as outlined in section 2.1, and (ii) using the same gold-standard seed points but with seed jittering activated and *N*_jitter_ = 40 random additional seeds per initial seed point. The corresponding inference and postprocessing parameters were selected through a manual search for effective configurations on a small number of training images. In both cases, the raw, unedited results were kept for evaluation. All evaluation metrics were then computed over the full collection of generated reconstructions.

We compare our model with all algorithms from the BigNeuron benchmarking study Manubens-Gil et al. (2023) for which reconstructions were available. For each algorithm, we computed all evaluation metrics described in 2.8 on the given reconstruction SWCs, using the corresponding gold standard SWCs (after topology correction, see 2.1) as the reference. Note that, although we provide as direct a comparison as possible, the reference reconstructions used to evaluate our method had the nodes inside the soma removed as detailed in section 2.1.

A description of the values we chose for every parameter in our algorithm is shown in the box below.

#### Final Parameters

**Environment parameters:** Target step length: 5.0 vox, Stop threshold: 1.0 vox,

**Training parameters:** Batch size: 64, Learning rate: 0.001, Update steps (*T*_update_): 25000, Number of rounds (*N*_rounds_): 50, Policy roll-in rounds (*T*_DAgg_): 25000, Warm-start steps (*T*_ws_): 100000, Roll-in probability start (*β*_start_): 0.95, Roll-in probability minimum (*β*_min_): 0.1, Roll-in probability step (*β*_step_): 0.05, Buffer capacity: 100000, *N*_jitter_: 0.

**Loss parameters:** Continue-norm threshold (*τ*): 1.0, Continue-direction weight (*λ*_dir_), Continue weight (*c*): 1.0, Classification weight (*λ*_cls_): 1.0, Classification temperature (*T*): 0.2

**Postprocessing parameters:** Merge threshold: 8.0, Confidence threshold: 2, Length filtering: None Resample step size: 1.0, Smoothing: None

**Evaluation parameters:** Tolerance (*τ*): 2.0

## 3. Results

The average symmetric geometric distance between our reconstructions and the gold standard across the full dataset was slightly smaller when seed jittering was activated (6.69 voxels) versus without (8.15 voxels), though both settings occupy the same rank when compared with all algorithms in geometric distance. For consistency, we show only the results with jittering activated. All results including those with and without seed jittering enabled are provided in Supplementary Table 1.

### 3.1. Qualitative Results

Figure 4 presents several reconstructions overlaid on their corresponding images. We highlight three examples with relatively strong geometric correspondence to the gold standard (Figure 4, A) and three with relatively poor correspondence (Figure 4, B). Visual inspection of the low-quality reconstructions clarifies the sources of error: in the left and middle panels, the traced neurites erroneously cross into and follow neighboring neurons, whereas in the right panel, a lower signal-to-noise ratio compared to most of the training set leads to spurious tracing of background structures. All erroneous reconstructions were subsequently corrected using the editing tools in our GUI, requiring only a small number of operations. As shown in Figure 4, C, the majority of corrections involved removing neighboring neurons that were incorrectly joined. While other groups have developed automatic postprocessing methods for this task Li et al. (2019), we found our semiautomatic approach to be sufficient for this dataset, as it only took about 30 seconds per neuron.

**Figure 4:**
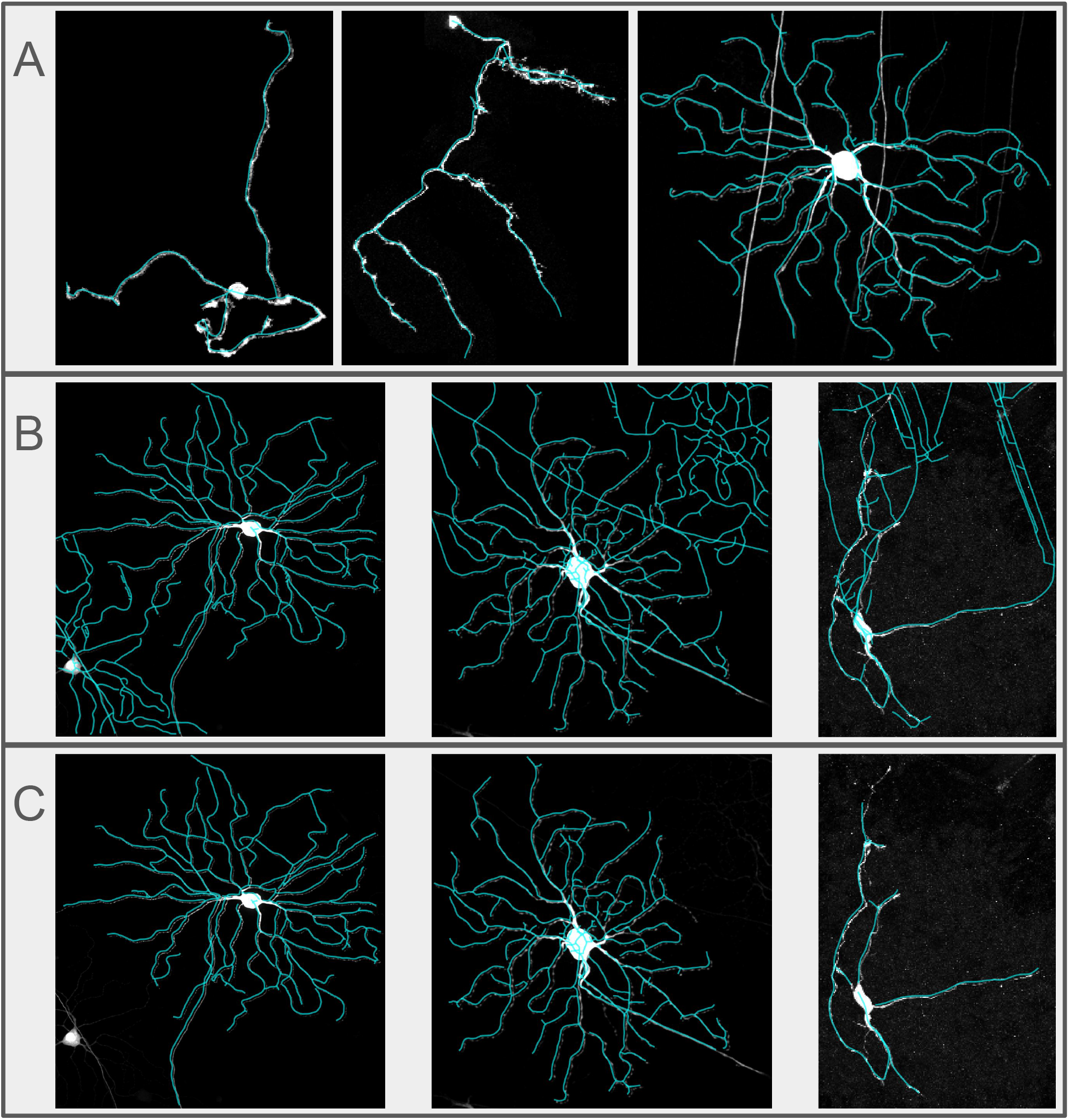
(A,B) Six examples of fully automated neuron reconstructions. (A) Examples with low symmetric geometric distance. (B) Examples with high symmetric geometric distance. (C) The result of manual correction for each high-distance trace in row B. The left and middle traces were fixed with just a few clicks by pruning branches at nodes whose descendants had incorrectly followed a neighboring neuron’s dendrites in the same image or extended into background regions. The right trace was repaired by rerunning the tracing with seed jittering turned off, which had previously caused spurious traces of background noise. We did not observe any neurons that were severely undertraced; however, if an initial pass misses substantial portions of a neuron, our manual interface enables the user to initiate new traces from additional seeds and then join these segments to the original reconstruction (see Figure 7 (D,E)), or to make fully manual corrections if preferred. Trace overlays are slightly displaced in the figure to improve the visibility of the neuron.

### 3.2. Quantitative metrics

Because each algorithm for comparison provided a different set of reconstructions (as provided by the the Gold166 publication Manubens-Gil et al. (2023)), we implemented a direct comparison between each pair of algorithms by computing the mean geometric distance for only the reconstructions shared between each pair. For each algorithm, we counted the number of match-ups for which it had the smaller mean distance. The proportion of winning match-ups is presented for every algorithm in Figure 5a. For a direct comparison with Manubens-Gil et al. (2023), we also compared algorithms using the mean geometric distance across all of their reconstructions (Figure 5b).

**Figure 5:**
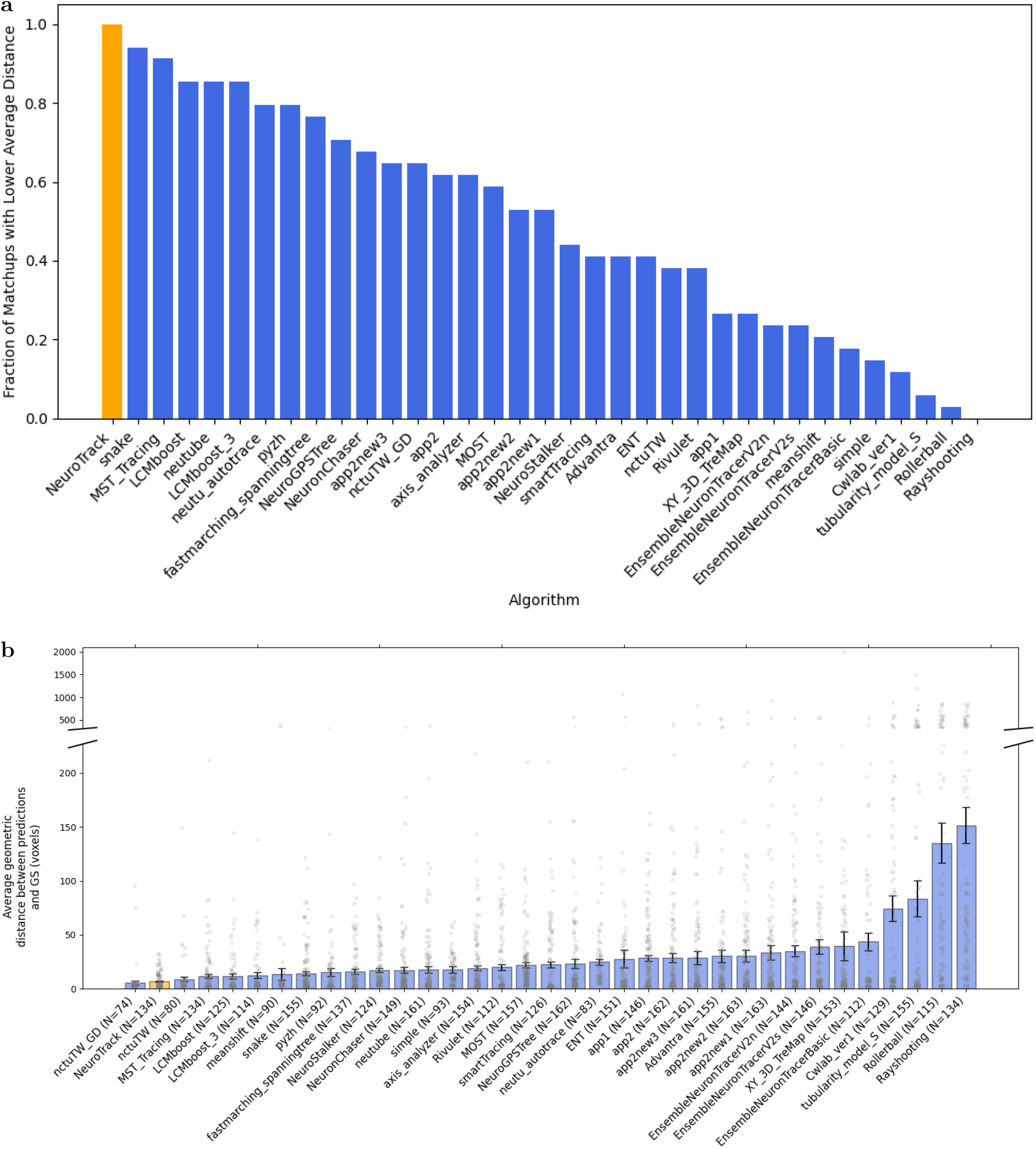
Quantitative results comparing the performance of different methods, shown as the proportion of winning direct pairwise matchups (a), and as the mean symmetric geometric distance across all neurons reconstructed by each algorithm (b).

### 3.3. Other accuracy measures

As in Manubens-Gil et al. (2023), we defined an aggregate similarity score by computing the normalized root mean square distance over all metrics (excluding precision and coverage since they are derived from the different structure fraction), after first scaling each metric to the range [0, 1], where 0 indicates a perfect match to the gold standard annotation and 1 corresponds to the largest discrepancy observed for that neuron across all algorithms. The mean aggregate similarity score is presented in Figure 6.

**Figure 6:**
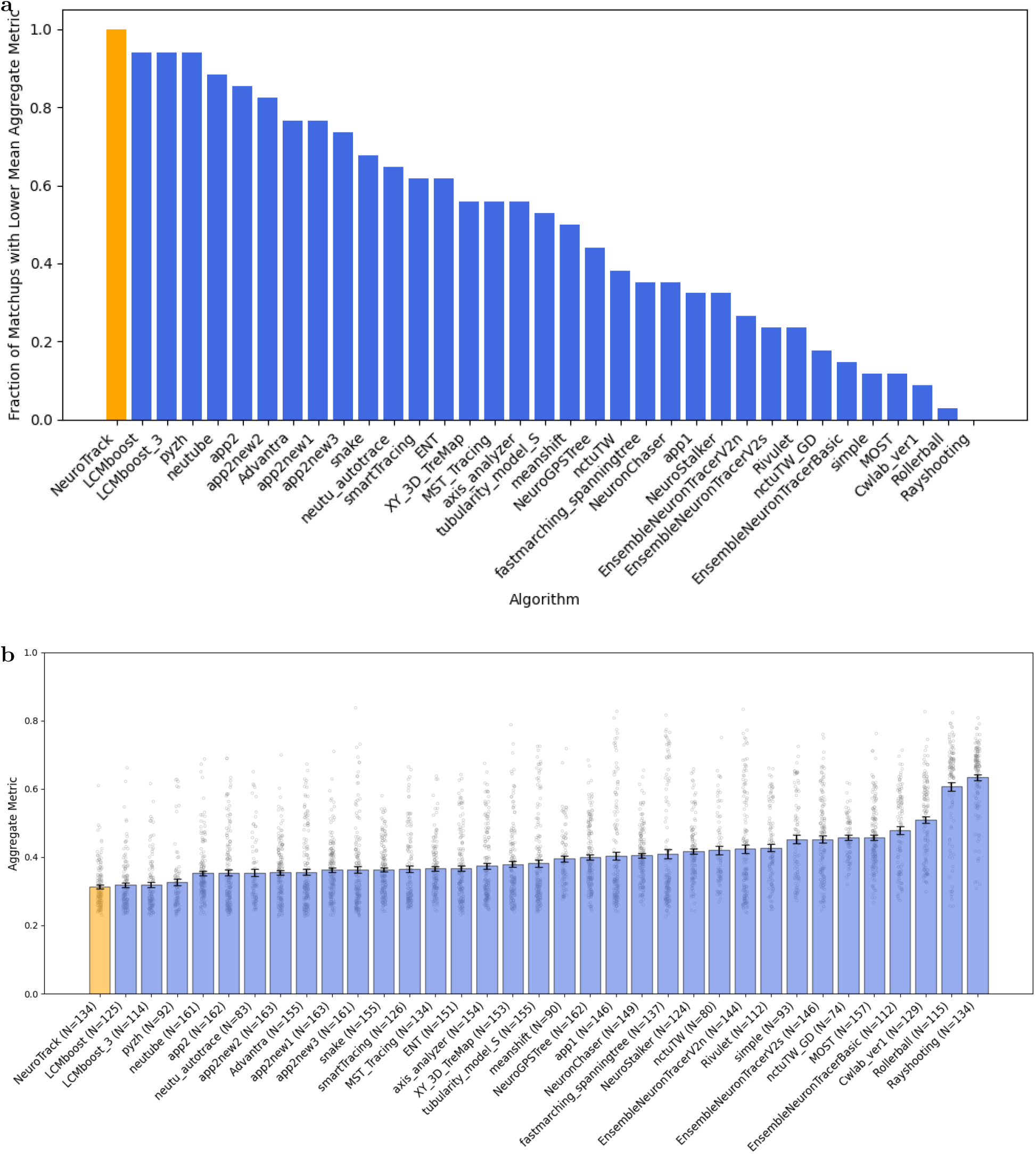
Quantitative comparison of the aggregate similarity score between each reconstruction and the gold standard, shown as the proportion of winning direct pairwise matchups (a), and as the mean value across all neurons reconstructed by each algorithm (b).

### 3.4. GUI Wrapper

We built a graphical tracing interface (GUI) designed as an interactive human-in-the-loop environment for neuron reconstruction, supporting the full workflow from seed placement through tracing, postprocessing, and quantitative evaluation. The interface presents synchronized orthogonal views of volumetric data (XY, XZ, YZ), enabling users to localize structures in 3D with maximum intensity projection and slice view options. In practice it is primarily used to curate seed points, launch model-based tracing for individual neurons or full image sets, and iteratively refine results by combining algorithmic inference with expert oversight. This design makes it suitable both for producing high-quality reconstructions and for diagnosing model behavior during development. This is in contrast to alternative software tools where AI techniques are provided as assistance to a largely manual process Zhang et al. (2024).

A key feature of the interface is its interactive analysis tools. Users can inspect prediction traces and reference annotations, edit traces through selection and clipping operations, adjust tracing seeds and set seed jittering parameters, and immediately re-run inference under updated settings. The same session also supports postprocessing controls (e.g., filtering, resampling, smoothing, and merge operations) and direct evaluation against ground-truth SWC data with report generation. By integrating navigation across image volumes, annotation editing, parameter management, and evaluation outputs in a single GUI, the system reduces context switching and enables reproducible expert-guided reconstruction workflows. An example of our GUI and its usage is shown in Figure 7.

**Figure 7:**
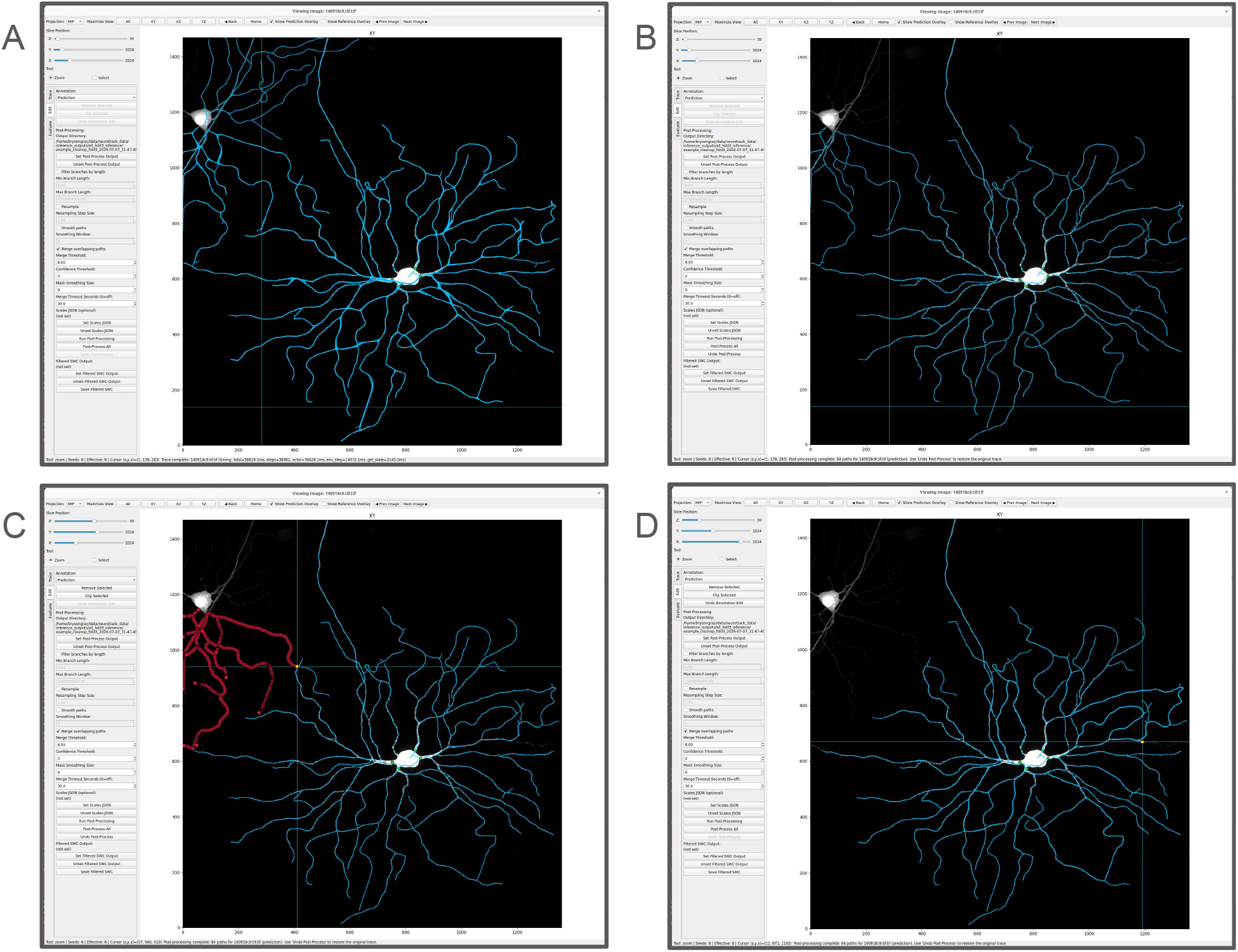
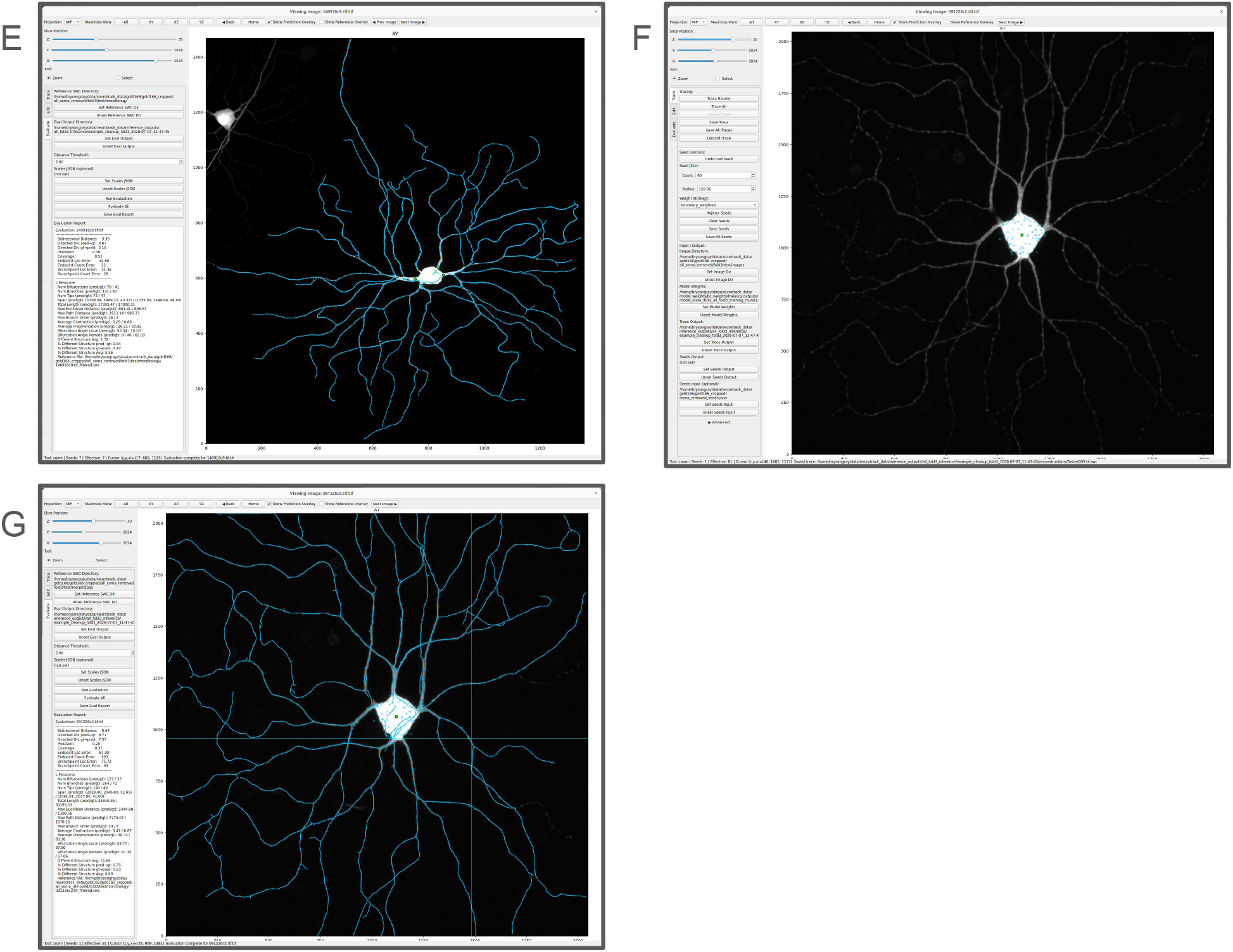
A) In the first step a user performs automated reconstruction. The resulting trace prior to preprocessing is displayed in blue over the neuron volume as a max intensity projection. B) The same reconstruction after performing merging with confidence clipping. C) This trace has mistakenly traced part of a neighboring neuron. The user can correct the error by selecting a node to prune. The downstream branches highlighted in red will be removed upon approval. D) A small segment of true neuron was removed during confidence filtering. This may also be remedied by selecting an existing node from which to begin a new trace. E) The corrected reconstruction after automated tracing from the selected node. F) Rather than placing seeds at the root of each primary dendrite, seed jittering allows the user to place one seed anywhere in the soma and allow the randomly jittered seeds to capture all the radiating dentrites. The jittered seeds can be seen as small blue dots, while the user selected point is shown as a green dot at the center of the soma. The resulting reconstruction is shown in panel (G).

## 4. Discussion

Our experimental results demonstrate that the proposed method is highly competitive with, and in the majority of cases (see Figures 5 and 6) superior to, 34 existing approaches for reconstructing tree-like structures. In pairwise, head-to-head comparisons based on the symmetric geometric distance metric, our method outperforms all other methods. It also ranks number one when aggregating all evaluation metrics. Although it ranks second overall in terms of mean symmetric geometric distance, it consistently outperforms those methods that were evaluated on the majority of images. This suggests that the method is both robust and reliable across a wide range of data, and that the small gap to the top-performing mean score is partly due to differences in evaluation coverage.

A key algorithmic property of our approach is that it guarantees tree-structured outputs from each seed by construction. Many existing methods may produce graphs with cycles or disconnected components, requiring additional postprocessing to enforce a tree topology. In contrast, our method directly encodes the tree constraint, which ensures that the final reconstruction is structurally consistent with the underlying anatomy and avoids ad hoc corrections.

Another practical advantage is that our method does not require a prior segmentation of the image as a preprocessing step. Many alternative algorithms rely on a binary or probabilistic segmentation to define a vessel or neurite mask, which introduces an additional potential source of error and typically demands substantial annotation or manual tuning. By operating directly on the image data without this prerequisite, our approach simplifies the pipeline and reduces dependence on upstream segmentation quality.

A central design choice of our algorithm is its sequential nature, which requires seed points. This can be viewed as a limitation in the sense that fully automatic reconstruction is not achieved: at least one initial seed point must be provided, and additional seeds may be needed to initiate new branches or disconnected regions. However, this same property is also an advantage from a practical, interactive perspective. It naturally enables a semi-automatic workflow in which a user can provide new seed points to guide the reconstruction, correct omissions, or focus on regions of particular interest. This makes the method well suited for expert-in-the-loop applications, where full automation is less critical than controllability and reliability. Developing strategies to automatically detect suitable seed points, or to integrate seed detection into the reconstruction itself, is an interesting direction for future work that could further reduce user intervention.

An additional computational limitation is that our method was designed for imaging datasets to fit in memory. Important contributions toward scaling up neuron segmentations tools, using remote compute and collaborative editing, have been made by other groups Zhang et al. (2024); Winnubst et al. (2019).

A further limitation concerns the computation and interpretation of the distance metrics used for comparison with the original benchmarking study Manubens-Gil et al. (2023). In the published results, distances were reported using the neuron distance plugin for Vaa3D. We re-evaluated the methods using both that plugin and our own implementation of the distance measure. However, we were unable to fully reproduce the distances reported in the original work. As a consequence, there is an inherent uncertainty in directly comparing our absolute distance values with those originally published. For consistency, we base our comparative conclusions on distances computed using our implementation. To ensure transparency, we provide the Vaa3D neuron distance plugin results for all reconstructions in Supplementary Table 2. Finally, the comparison with other algorithms is not perfectly one-to-one, because in our reference annotations we deleted nodes inside the soma to remove labels in areas where the correct step direction is ambiguous and therefore not learnable by the model. Nevertheless, we expect that the removed nodes constitute only a very small portion of the overall neuron and thus introduce only a minor discrepancy.

In summary, our method achieves strong performance across evaluation metrics, guarantees tree-structured outputs, and removes the need for segmentation as a preprocessing step. Rather than being a constraint, the use of seed points enables a practical expert-in-the-loop workflow: users can rapidly initialize tracing, focus effort on difficult regions, and iteratively correct any errors with direct control. Developing automated seed selection methods and adapting this tool for analysis of large scale datasets are promising avenues for future research.

## Acknowledgements

This work was supported by The National Institutes of health through grants R01 NS121761, UM1 NS132173.

## Data availability

All data required to reproduce the results of this study have been deposited on figshare under the DOI: https://doi.org/10.6084/m9.figshare.33113558. The repository includes: neuron reconstructions generated by the proposed method; the soma-removed gold-standard reconstructions used for training and testing; the cleaned gold-standard reconstructions used to evaluate alternative methods; the cropped TIFF image volumes used for training and testing; tracing seed points; and model weights for all five training folds.

## Code Availability

The custom software developed for this study is open source and available in the GitHub repository https://github.com/BrysonGray/neurotrack.git. The exact version of the source code used for the peer-reviewed manuscript is permanently archived on figshare under the DOI: https://doi.org/10.6084/m9.figshare.33116063. The trained model weights are archived on the Hugging Face Hub at https://huggingface.co/brysongray/NeuroTrack under the DOI: https://doi.org/10.57967/hf/9682.

