## Supplemental for "Automated Neuron Tracing with Imitation Learning"

**Supplementary Information for "Automated Neuron Tracing with Imitation Learning" by Bryson Gray and Daniel Tward, 2026**

**Contents**

**Supplementary Information** **1**

**1 Supplementary Methods** **2**

1.1 Dataset filtering and preprocessing . . . . . . . . . . . . . . . . . . . . . . . . . . . . . . . . . . 2

**2 Supplementary Figures** **2**

**3 Supplementary Tables** **4**

**4 Source Data** **4**

### 1. Supplementary Methods

#### *1.1. Dataset filtering and preprocessing*

Several subsets of the BigNeuron Gold166 challenge dataset were excluded because their corresponding gold-standard reconstructions did not reliably follow the neurite centerlines evident in the image data and could not be corrected by applying a uniform spatial offset. Additional volumes were excluded because they contained binary intensity values rather than raw image intensities, resulting in insufficient contrast in densely arborized neurite regions. The excluded datasets were: `chick_uw`, `zebrafish_larve_RGC_UW`, `mouse_korea`, `mouse_ugoettingen` (don't reliably follow the neurite centerlines), and `silkmoth_utokyo` (binary intensity values).

Many of the retained gold-standard reconstructions required additional preprocessing to be suitable for use as training targets. The main issues and corresponding corrections were as follows:

1. **Duplicate node identifiers**. Some SWC files contained multiple nodes sharing the same identifier (ID). To resolve this, we detected all duplicates and retained only the node whose parent index differed from its own ID by one (i.e., a consistent local indexing relationship), while marking all other nodes with the same ID for removal.
2. **Nodes without parents (multiple roots)**. Numerous files contained more than one node with no parent (parent ID = $-1$). We designated the first such node as the root. For each additional parentless node, we searched for the nearest node belonging to a different connected component that already had a valid parent and, if this candidate parent lay within a Euclidean distance of 10 units, we reassigned the parentless node's parent to this nearest node. If no suitable nearby parent was found, the no-parent node was removed. In cases where the nearest non-connected node occupied exactly the same coordinates as the no-parent node, we removed the parentless node and reassigned its children to the node at the shared coordinates.
3. **End nodes with shared coordinates across components**. We additionally inspected terminal nodes (degree-1 nodes) that shared coordinates with other nodes. When such nodes belonged to distinct connected components, we merged the components by connecting them and designating the component with the larger number of nodes as the parent subtree. To maintain a consistent tree structure, we reversed the parent–child relationships throughout the smaller component so that it attached correctly to the larger one.

Following these cleaning operations, we validated the integrity of each SWC reconstruction by re-checking for nodes without parents, duplicate node IDs, multiple disconnected components, and the presence of cycles, ensuring that each file represented a valid rooted tree (or arborization) suitable for downstream analysis.

For the `taiwan_flycircuit` subset of the data, we observed a systematic spatial offset: all node coordinates were shifted by $+1$ unit along each spatial axis relative to the underlying image volumes. We corrected this alignment error by uniformly subtracting 1 unit from all x, y, and z coordinates in the corresponding SWC files.

### 2. Supplementary Figures

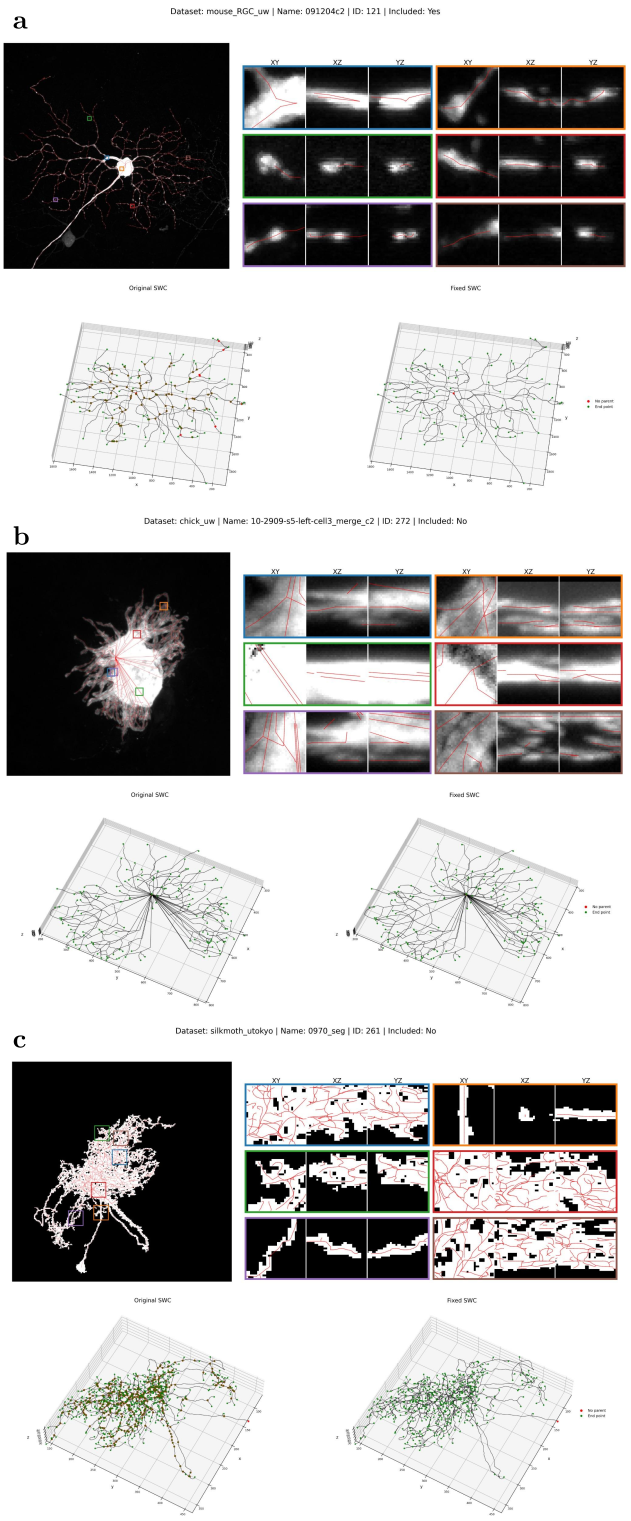


Figure S1: Data quality visualization for data selection. Each panel shows the full image and gold standard SWC as a maximum intensity projection in the XY plane (top left), Expanded views of six image patches randomly selected along the neuron skeleton projected along each image axis with the in-frame skeleton overlaid (top right). The original SWC skeleton is shown (bottom left) next to the corrected skeleton (bottom right) with nodes with no parent shown in red and end points shown in green. a) A neuron that was included in the training dataset. b) A neuron that was excluded from the training dataset because much of the neuron's structure has ambiguous directionality or the gold standard skeleton is off-center from the perceived center-line. c) A neuron that was excluded because the image has binary intensity values, making neurites undifferentiated in densely arborized regions

### 3. Supplementary Tables

This section contains a description of our supplementary tables, which are included in a separate spreadsheet file.

**Supplementary Table 1**
Evaluation results on all metrics proposed between reconstructions and gold standard for each neuron reconstruction for all algorithms.

**Supplementary Table 2**
Vaa3D neuron_distance plugin outputs for distances between reconstructions and gold standard for each neuron reconstruction for all algorithms.

**Supplementary Table 3**
Lookup table matching each neuron name and data subset to its ID number along with its inclusion status in the training data.

**Supplementary Table 4**
Each Neuron ID along with the cross-validation test fold (1-5) in which it was included.

### 4. Source Data

This section contains a description of our figure source data, which is included in a separate spreadsheet file.

**Source Data Fig. 5a**
Table showing the proportion of winning direct pairwise matchups between algorithms. For each pair of algorithms, the one with the lower symmetric geometric distance for its reconstructions, including only the neurons that both algorithms provided reconstructions for is counted as winning.

**Source Data Fig. 5b**
Mean and median geometric distances between each algorithm's reconstructions and gold standard annotations and the number of reconstructions each algorithm created.

**Source Data Fig. 6a**
Table showing the proportion of winning direct pairwise matchups between algorithms. For each pair of algorithms, the one with the lower aggregate similarity score for its reconstructions, including only the neurons that both algorithms provided reconstructions for is counted as winning.

**Source Data Fig. 6b**
Mean and median aggregate similarity scores between each algorithm's reconstructions and gold standard annotations and the number of reconstructions each algorithm created.
